# Deciphering the Pathogenic Landscape of Amyloid Light-Chain Cardiomyopathy

**DOI:** 10.1101/2025.03.19.644248

**Authors:** Qunchao Ma, Zhuo Wang, Jinyun Zhu, DanDan Yang, Xiaohua liu, Jichen Xu, Xiaohong Pan, Ning Zhang

## Abstract

Amyloid light-chain cardiomyopathy (ALCA) is an infiltrative disorder marked by misfolded immunoglobulin light-chain deposition in the myocardium, ultimately leading to cardiac dysfunction. Despite its clinical severity, the underlying mechanisms remain poorly understood. Here, we integrated multi-omics analyses of human cardiac samples to construct a comprehensive cellular and spatial atlas of the ALCA heart.

We observed a marked expansion of PTX3^+^ fibroblasts (FB), which undergo a distinct phenotypic transition into pro-fibrotic POSTN^+^ FB regulated by EGR1.Concurrently, SPP1^+^ macrophages (Mac) emerged as major drivers of fibrosis, interacting robustly with PTX3^+^ FB via APP–CD74, GAS6–MERTK, and TGF-β1–TGF-β1/2 pathways. Endothelial cell (EC) profiling revealed substantial vascular remodeling characterized by the emergence of specialized capillary-like immune endothelial cells expressing chemokines CXCL1, CXCL3, and CCL2, alongside depletion of functional capillary EC. Immunologically, elevated cytotoxic CD8^+^ T cells and reduced NK cells contributed to an imbalanced inflammatory milieu, with NFκB2 orchestrating both fibrotic and immune pathways across multiple cell types.

These findings highlight the pivotal role of fibroblast–immune crosstalk, particularly the SPP1^+^ Mac–PTX3^+^ FB axis, in driving ALCA pathogenesis. Targeting these pathological cellular interactions may offer a promising therapeutic avenue to mitigate fibrosis, restore immune homeostasis, and improve cardiac function in ALCA.

## Introduction

Cardiac amyloidosis (CA) is an infiltrative cardiomyopathy characterized by the deposition of insoluble amyloid fibrils, ultimately leading to cardiac dysfunction^1^. Among its subtypes, light-chain cardiac amyloidosis (ALCA) is particularly aggressive, resulting from the deposition of immunoglobulin light chain fibrils and often associated with poor survival outcomes^2,3^. Clinically, ALCA typically presents as a restrictive cardiomyopathy, with an estimated incidence of 8–12 cases per million and a mortality rate of 0.58 per 1,000 cases annually^4^. Without timely intervention, the median survival after diagnosis is less than one year^5^, underscoring the urgent need for improved therapeutic strategies^2^.

The pathophysiology of ALCA is multifactorial, encompassing both the proteotoxic effects of misfolded light chains and the mechanical disruption caused by amyloid fibril deposition^6^. Although recent studies have clarified aspects of this toxicity, comprehensive insights into the cellular and molecular interactions underlying cardiac injury in ALCA remain limited. Additionally, the lack of representative animal models complicates translating research insights into clinical therapies.

Emerging evidence emphasizes the importance of cellular heterogeneity in cardiac tissues, demonstrating that distinct populations of cardiomyocytes, fibroblasts, immune cells, and endothelial cells contribute differently to disease progression^7,8^. Moreover, spatial organization of these cell types critically influences pathogenic processes, including inflammation, extracellular matrix (ECM) remodeling, and vascular dysfunction^9–12^. Despite significant progress made in other cardiomyopathies^9,13,14^, similar investigations remain sparse in the context of ALCA.

In this study, we provide a detailed analysis of cellular composition, spatial relationships, and molecular interactions in ALCA hearts. By elucidating these intricate interactions, we aim to uncover novel therapeutic targets and pathogenic mechanisms central to ALCA progression.

## Results

### Overall Cellular Landscape of ALCA

To investigate the cellular composition and heterogeneity underlying ALCA, we analyzed cardiac tissue samples from both ALCA patients and healthy groups (HG) using imaging mass cytometry (IMC) and single-cell RNA sequencing (scRNA-seq) (Fig. 1A). Histological staining with hematoxylin-eosin (HE), Masson’s trichrome, and Congo Red, along with lineage-specific IMC markers, revealed marked disruption of myocardial architecture in ALCA, especially in fibrotic and inflammatory regions (Fig. 1B). IMC analysis identified fibroblasts (FBs), endothelial cells (ECs), and macrophages as the predominant cell types, alongside an undefined population likely corresponding to cardiomyocytes (Fig. 1C, Supplementary Fig. 1-2). ScRNA-seq further identified FB, EC, myeloid cells, proliferating cells, smooth muscle cells, and various lymphocyte subsets (T, B, and NK cells), illustrating the complex cellular environment in ALCA (Fig. 1D–E, Supplementary Fig. 3).

**Fig. 1.**
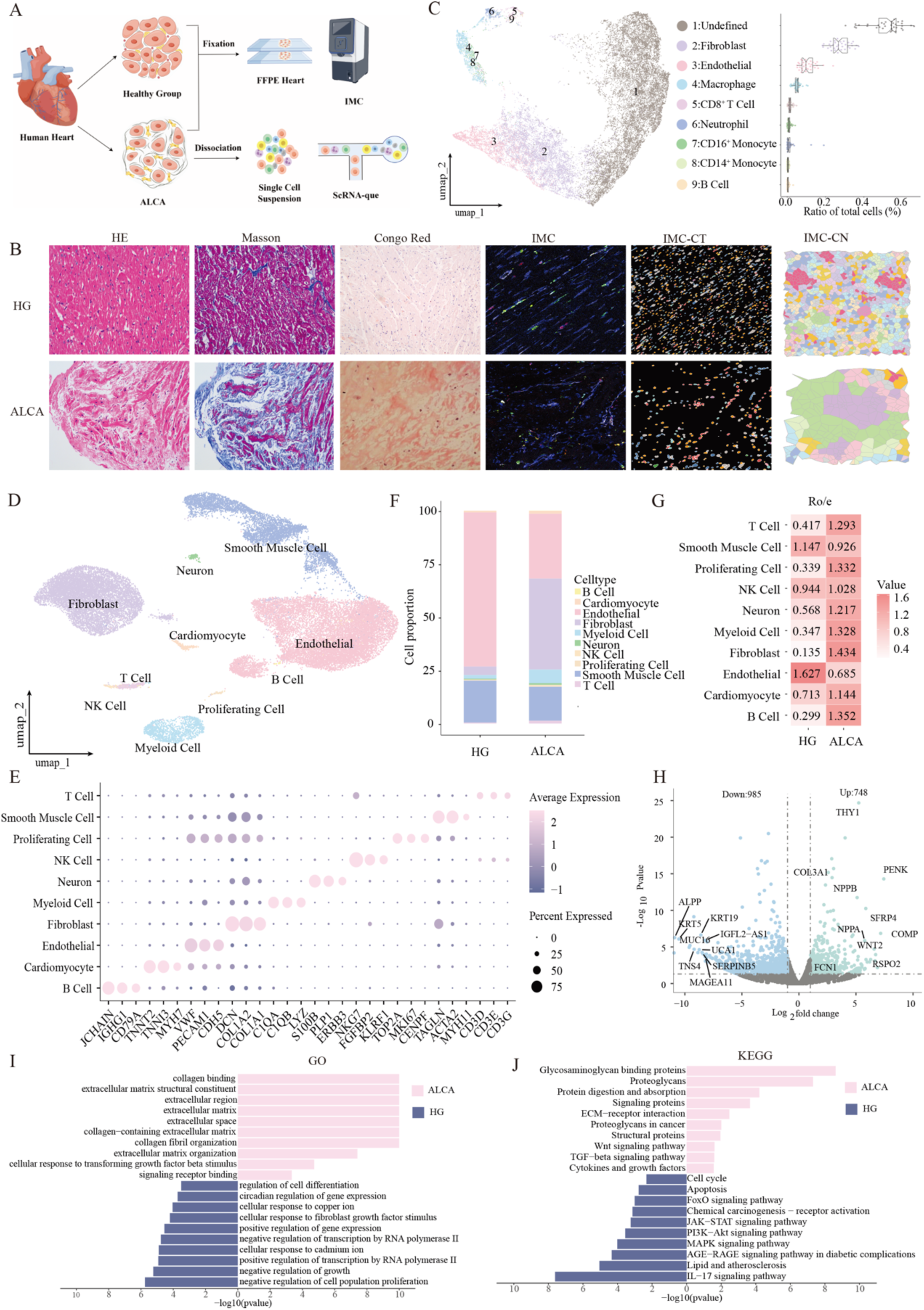
Study Overview and Cell Subset Identification. **(A)** Graphic summary of the study design showing single-cell RNA sequencing (scRNA-seq) for ALCA hearts and imaging mass cytometry (IMC) for both healthy (HG) and ALCA tissues. **(B)** Representative images of pathological staining in HG and ALCA hearts, including Hematoxylin and eosin (HE) images (left panel, first column), Masson’s trichrome staining (left panel, second column), Congo Red staining (left panel, third column), IMC staining (right panel, third column), cell type segmentation of IMC staining (right panel, second column), cellular neighborhoods of IMC staining (right panel, first column). **(C)** UMAP of IMC data showing major cell types, with a box plot summarizing cell counts across nine cell types. **(D)** UMAP of scRNA-seq data depicting cell clusters. **(E)** Dot plot of canonical marker expression in each cluster. **(F)** Stacked bar chart of scRNA-seq cell-type proportions. **(G)** Ro/e scores showing tissue prevalence of each cluster. **(H)** Volcano plot depicting differentially expressed genes (DEGs) from RNA-Seq analysis. Blue dots represent significantly downregulated genes, while green dots represent significantly upregulated genes. **(I)** The bar graph illustrates Gene Ontology (GO) enrichment analysis derived from RNA-Seq DEGs. GO terms are grouped according to biological processes, with the bar height representing the degree of enrichment. **(J)** The bar graph presents Kyoto Encyclopedia of Genes and Genomes (KEGG) pathway enrichment analysis from RNA-Seq DEGs. Pathways are ranked based on their significance, with the bar height indicating the enrichment level.

Notably, FB and myeloid cells were significantly expanded in ALCA, whereas EC populations were decreased, reflecting a structural shift driven by fibrotic remodeling (Fig. 1F-G). Enrichment ratios for FBs and myeloid cells reached 1.434 and 1.328, respectively, while ECs declined to 0.713. Differential gene expression analysis revealed upregulation of pathways related to the ECM organization, TNF signaling, and immune responses, highlighting the intricate remodeling processes that characterize ALCA (Fig. 1H-J).

### Fibroblast Heterogeneity in ALCA

Among the observed cellular changes, FB expansion was particularly prominent in ALCA compared to HG (Fig. 2A). These FB displayed elevated expression of ECM-related cytokines (POSTN, FAP, COL1A1) and genes associated with TGF-β receptor signaling (NREP, FOSB), indicating a pro-fibrotic phenotype (Fig. 2B-C, Supplementary Fig. 4A). Clustering analysis revealed eight distinct FB subtypes, including PTX3^+^ FB, POSTN^+^ FB, SCN7A^+^ FB, FABP4^+^ FB, myofibroblasts (MyoFB), antigen-presenting fibroblasts (AP FB), PCOLCE2^+^ FB, and endothelial-like antigen-presenting fibroblasts (Endo-like AP FB) (Fig. 2D-E). PTX3^+^ FB and POSTN^+^ FB were the predominant subtypes in ALCA, whereas SCN7A^+^ FB was more abundant in HG (Fig. 2F, Supplementary Fig. 4B-C).

**Fig. 2.**
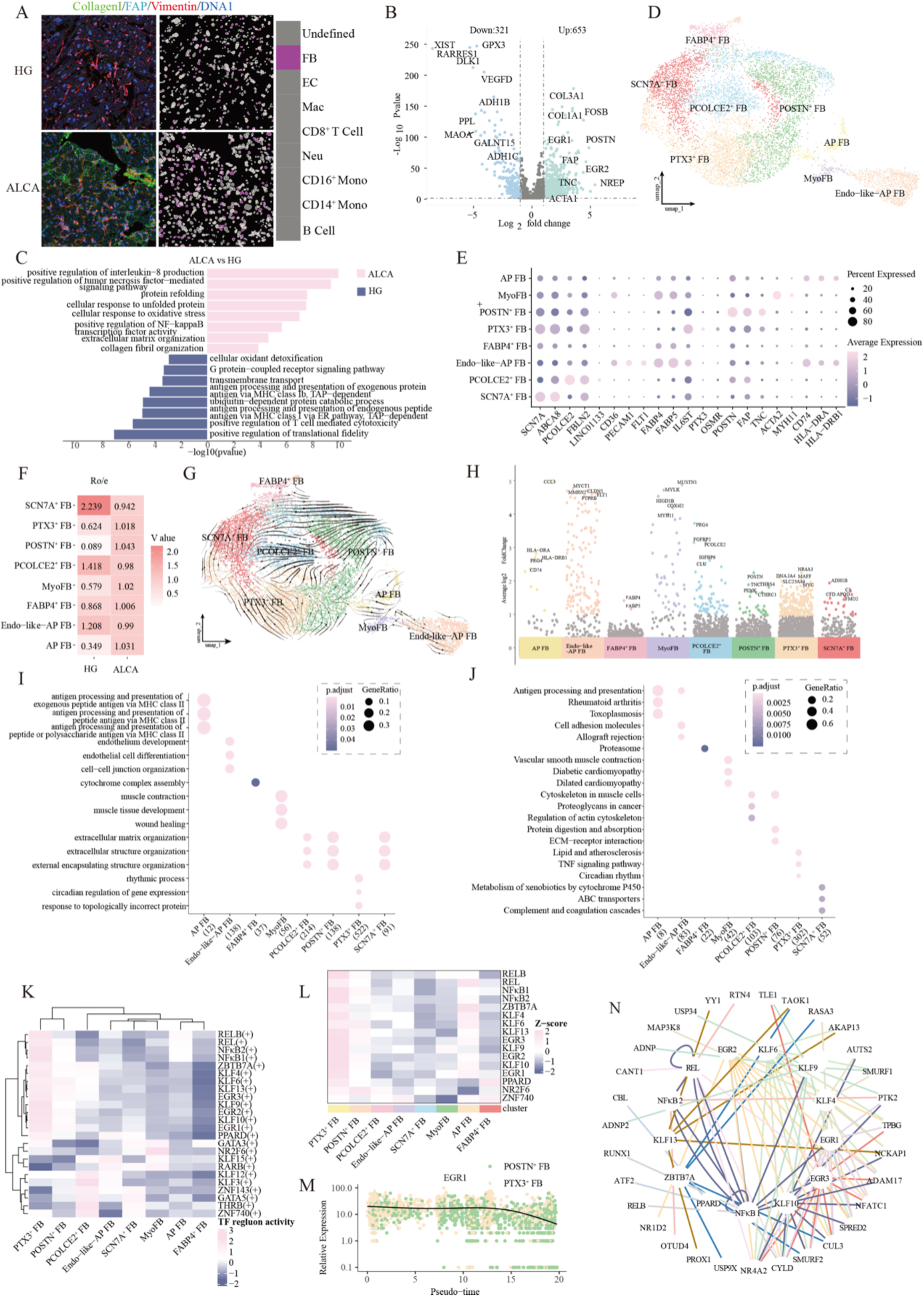
Characterization of fibroblast in ALCA. **(A)** IMC images (left panel) and single-cell masks(right panel) showing fibroblast markers, including Collagen I (lime), FAP (cyan), Vimentin (red) and DNA1 (blue) in HG and ALCA hearts. **(B)** Volcano plot of DEGs in fibroblast clusters from scRNA-seq. Blue dots represent significantly downregulated genes, while green dots indicate significantly upregulated genes. The x-axis displays the log2 fold change, and the y-axis represents the -log10(p-value). **(C)** Bar plot of GO enrichment for fibroblast DEGs by single-cell RNA sequencing. The x-axis represents the -log10(p-value), and the y-axis lists the significantly enriched GO terms categorized into biological processes. **(D)** UMAP displaying eight FB subsets. **(E)** Dot plot of marker expression across FB clusters. **(F)** Ro/e scores of FB subsets. **(G)** Velocity streamlines in a UMAP embedding showing the differentiation trajectory of FB clusters. **(H)** Top five marker genes for each FB subtype, highlighting key drivers of fibroblast identity. **(I)** GO dot plots for the FB subclusters. **(J)** KEGG dot plots for the FB subclusters. **(K)** Heatmaps of TF motif activity values, with the top five motifs per FB clusters. **(L)** Heatmaps of TF expression values for FB clusters. **(M)** EGR1 expression over pseudotime in POSTN+ FB (green) and PTX3+ FB (yellow)(top). Gene expression density plot for EGR1 (bottom). **(N)** Predicted FB signature genes (outer circle) and their corresponding TF regulons (inner circle).

Trajectory analysis suggested a differentiation trajectory from PTX3^+^ FB to POSTN^+^ FB, representing a progression from early activation to advanced fibrotic states (Fig. 2G, Supplementary Fig. 4D). PTX3^+^ FB expressed genes such as NR4A3, DNAJA4, MAFF (Fig. 2H), implicated in immune regulation and fibrosis^15–17^, and were enriched in circadian regulation, response to misfolded proteins, and lipid metabolism (Fig. 2I). These findings point to a multifaceted role for PTX3^+^ FB in initiating fibrosis, potentially influenced by circadian rhythms and lipid homeostasis. POSTN^+^ FB strongly expressed POSTN, TNC, and THBS4 (Fig. 2H), emphasizing their role in late-stage fibrosis through ECM deposition and structural remodeling^18,19^.

Pathway enrichment analysis further indicated that both PTX3^+^ and POSTN^+^ FB engage TGF-β signaling, TNF signaling, and ECM–receptor interactions (Fig. 2H-J). Several critical transcription factor (TF) regulons such as EGR, KLF9, NFκB1/2, REL, were shared by these FB subtypes, driving key fibrotic targets such as COL3A1, COL1A1, and FAP (Fig. 2K-L). EGR1 emerged as particularly active in PTX3^+^ FB, where it targets pro-fibrosis genes, suggesting that it may regulate the transition from early-stage (PTX3^+^) to advanced (POSTN^+^) fibrosis (Fig. 2M-N, Supplementary Fig. 4E-F).

The loss of SCN7A^+^ FB, which express genes such as ADH1B, FMO2, APOD, associated with ECM homeostasis and adipose differentiation^20,21^, exacerbate fibrotic pathology in ALCA by removing a regulatory influence on ECM and lipid balance (Fig. 2H, I and J).

### Endothelial Cell Heterogeneity in ALCA

A significant reduction in EC populations was observed in ALCA compared to HG (Fig. 3A), consistent with the known role of endothelial dysfunction in ALCA pathogenesis^22–24^. ECs in ALCA exhibited elevated levels of endothelial dysfunction markers (PTN, SERPINF1) and fibrosis-related genes (LUM, COL3A1, COL1A1, DCN), implicating TNF signaling, oxidative stress, and ECM organization in disease progression (Fig.3B-C, Supplementary Fig. 5A). Clustering analysis identified six distinct EC subtypes: arterial EC, venous EC, capillary EC, capillary-like immune EC, fibroblast-like EC (FB-like EC), and smooth muscle cell-like EC (SMC-like EC) (Fig. 3D–E). Compared with HG, ALCA showed a marked expansion of FB-like EC, venous EC, and capillary-like immune EC, accompanied by depletion of capillary EC and SMC-like EC (Fig.3F, Supplementary Fig. 5B-C), suggesting disrupted vascular homeostasis and endothelial plasticity. Trajectory analysis suggested that capillary EC can differentiate into other subtypes (Fig. 3G, Supplementary Fig. 5D). FB-like EC expressed high levels of fibroblast-associated genes (COL14A1, PTN, OGN), aligning with a fibrotic profile. Venous EC exhibited enhanced expression of genes (ACKR, SELP, PLAT, POSTN) linked to adhesion and remodeling. Capillary-like immune EC upregulated RND1, CXCL1, CXCL3, and CCL2, indicating a role in immune responses and antigen presentation (Fig. 3H).

**Fig. 3.**
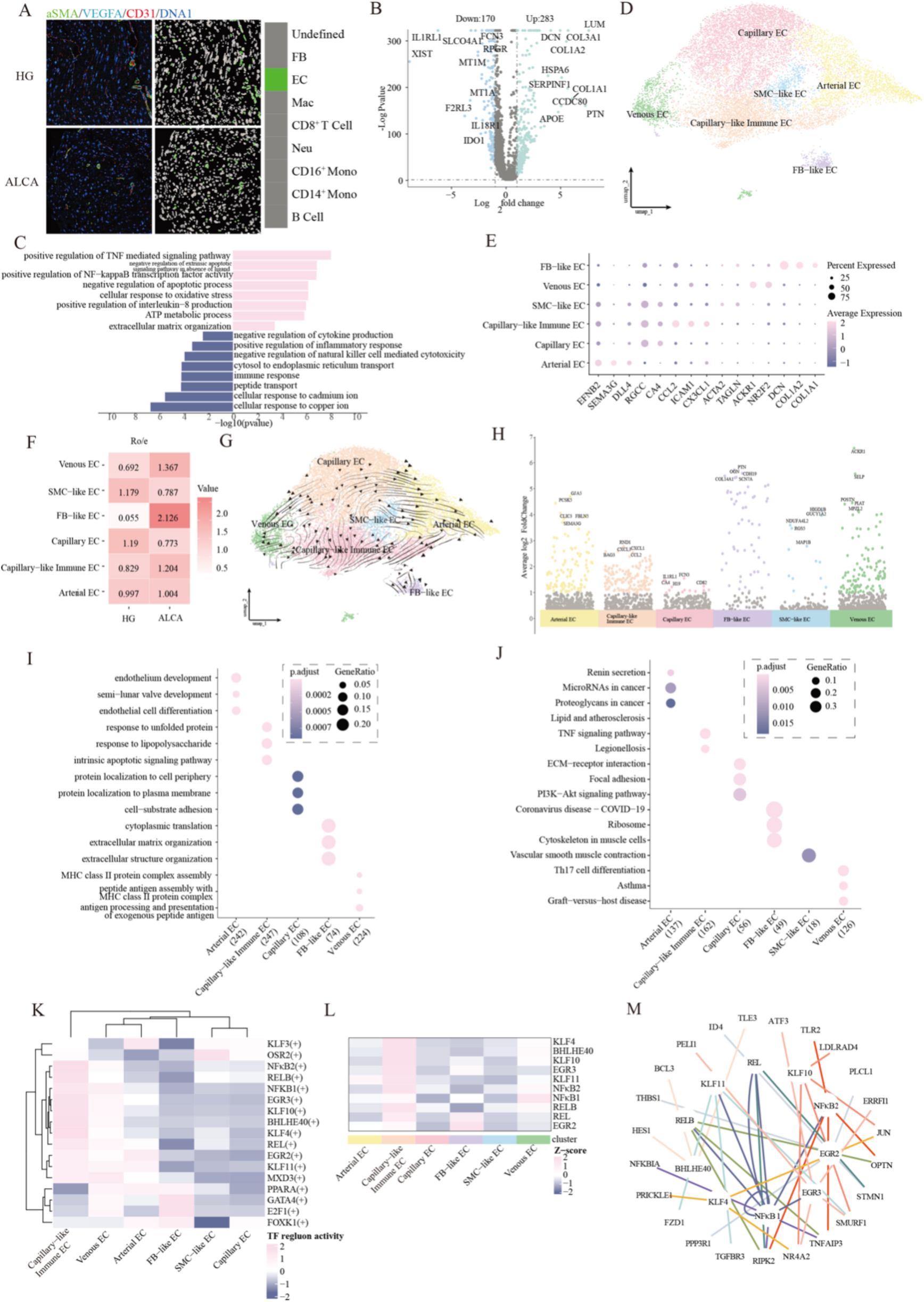
Characterization of endothelial cell in ALCA. **(A)** IMC images (left panel) and cell-type masks (right panel) showing EC markers, including αSMA (lime), VEGFA (cyan), CD31 (red) and DNA1 (blue) in HG and ALCA hearts. **(B)** Volcano plot of DEGs in EC clusters by scRNA sequencing. Blue dots represent significantly downregulated genes, while green dots indicate significantly upregulated genes. **(C)** Bar plot of GO enrichment for EC DEGs. **(D)** UMAP with six EC subsets identified by scRNA-seq. **(E)** Dot plot of key marker expression in each EC cluster. **(F)** Ro/e scores displaying EC subtype prevalence. **(G)** Velocity streamlines indicating EC differentiation trajectories. **(H)** Top marker genes for each EC subtypes. **(I)** GO dot plots for the EC subclusters. **(J)** KEGG dot plots for the EC subclusters. **(K)** Heatmaps of TF motif activity values, with the top five motifs per EC clusters. **(L)** Heatmap of TF expression values for EC clusters. **(M)** Circos diagram linking EC signature genes (outer circle) to their TF regulons (inner circle).

Functional enrichment analyses revealed that venous EC and capillary-like immune EC contribute to key immune-related processes, including responses to unfolded proteins, antigen presentation, and ECM organization (Fig. 3I). KEGG enrichment corroborated these findings by identifying TNF signaling, Th17 cell differentiation, and cell adhesion as critical pathways (Fig. 3J). These processes are governed by TFs such as NFκB1/2, EGR, and KLF, reflecting the pro-inflammatory phenotype of EC subtypes in ALCA. Notably, Bhlhe40, a TF involved in immunity under inflammatory conditions^25^, was significantly upregulated in capillary-like immune ECs, indicating its potential involvement in the inflammatory cascade of ALCA (Fig. 3K–M).

The depletion of capillary EC and SMC-like EC, coupled with diminished activity in pathways related to vascular integrity^26^, including endothelium development and protein localization to the plasma membrane (Fig. 3I-J), likely exacerbates vascular dysfunction in ALCA.

### Myeloid Cell Heterogeneity in ALCA

A marked infiltration of myeloid cells was observed in ALCA tissues compared with HG (Fig. 4A), underscoring the importance of myeloid-driven inflammation in disease progression. These cells showed elevated expression of genes related to inflammatory responses, G-protein-coupled receptor signaling, ECM organization, and chemokine pathways (Fig. 4B-C, Supplementary Fig. 6A). Clustering analysis identified thirteen myeloid cell subtypes, including monocytes (CD14^+^ Mono and CD16^+^ Mono), macrophages (SPP1^+^ Mac, MKI67^+^ Mac, CCL3^+^ Mac, LYVE1^+^ Mac, C1QC^+^ Mac, SMC-Mac, Endo-like Mac and FB-like Mac), neutrophils (Neu), mast cell and dendritic cells (CD1C^+^ cDC2) (Fig. 4D–E). Notably, FB-like Mac was unique to ALCA, while CCL3^+^ Mac and SPP1^+^ Mac were significantly expanded (Fig. 4F, Supplementary Fig. 6B-C), implicating them in fibrosis and inflammation^27,28^.

**Fig. 4.**
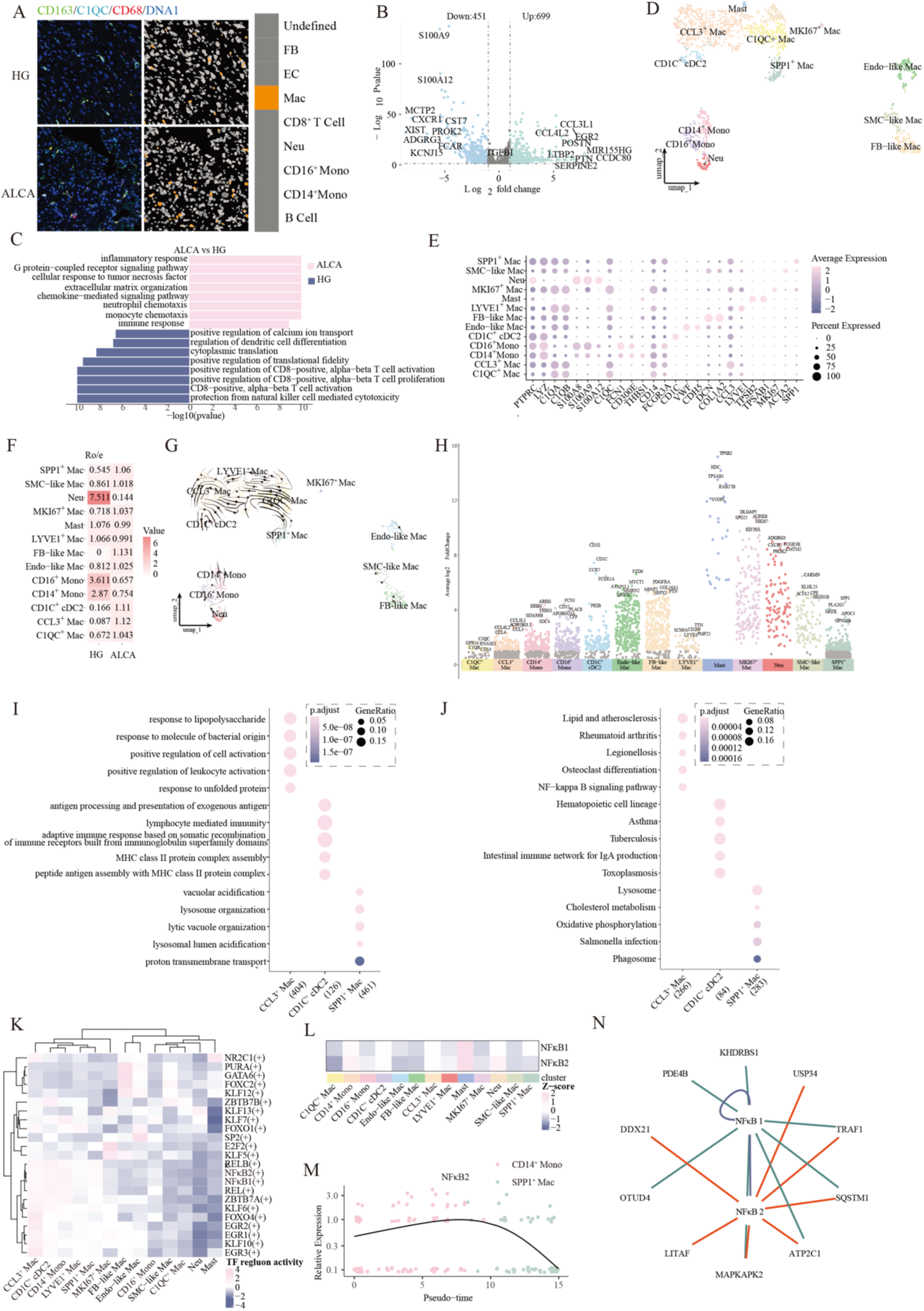
Characterization of myeloid cell in ALCA. **(A)** IMC images (left panel) and cell-type masks (right panel) with myeloid markers, including CD163 (lime), C1QC (cyan), CD68 (red) and DNA1 (blue) in HG and ALCA hearts. **(B)** Volcano plot of DEGs in myeloid clusters. Blue dots represent significantly downregulated genes, while green dots indicate significantly upregulated genes. **(C)** Bar plot of GO enrichment for myeloid DEGs. **(D)** UMAP showing thirteen myeloid cell subsets by scRNA-seq. **(E)** Dot plot of canonical markers for each myeloid cluster. **(F)** Ro/e scores indicating cluster prevalence. **(G)** Velocity streamlines of myeloid differentiation trajectories. **(H)** Top marker genes for each myeloid cell types. **(I)** GO dot plots for three prominent myeloid subtypes (SPP1^+^ Mac, CCL3^+^ Mac, CD1C^+^ cDC2). **(J)** KEGG dot plots for three prominent myeloid subtypes (SPP1^+^ Mac, CCL3^+^ Mac, CD1C^+^ cDC2). **(K)** Heatmaps of TF motif activity values, with the top five motifs per myeloid clusters. **(L)** Heatmap of TF expression values for myeloid subclusters. **(M)** NFκB2 gene expression over pseudotime in CD14^+^ Mono (pink) and SPP1^+^ Mac(green) (top). Gene expression density plot for NFΚB2 (bottom). **(N)** Circos diagram linking myeloid signature genes (outer circle) to TF regulons (inner circle).

Trajectory analysis revealed a progression from monocytes to SPP1^+^ Mac as the terminal differentiation state in ALCA (Fig. 4G, Supplementary Fig. 6D), implying that macrophage differentiation amplifies pro-inflammatory and pro-fibrotic signals. CCL3^+^ Mac expressed high levels of pro-inflammatory mediators (CCL3, CCL3L1, and CCL4L2), whereas SPP1^+^ Mac displayed elevated SPP1, PLA2G7, and APOC1 (Fig. 4H), linking them to both inflammation and lipid metabolism^29,30^.

Functional enrichment analyses revealed distinct roles for macrophage subtypes in ALCA. SPP1^+^ Mac contributed to vacuolar acidification, lysosome organization, and myeloid leukocyte activation, processes central to immune homeostasis, whereas FB-like Mac appeared to support structural remodeling by regulating ECM-related genes. CCL3^+^ Mac was associated with cytokine production and leukocyte recruitment, perpetuating chronic inflammation (Fig. 4I, Supplementary Fig. 6E). Pathway analysis further confirmed the involvement of lysosomal function, oxidative phosphorylation, and NFκB signaling (Fig. 4J, Supplementary Fig. 6E). Multiple TF regulons (RELB, NFκB1/2, REL) may drive the differentiation of CD14^+^ Mono into CCL3^+^ Mac and SPP1^+^ Mac (Fig. 4K-L, Supplementary Fig. 6F-G). By modulating targets in both myeloid cells (MAPKAPK2, DDX21) and fibroblasts (FAP, LRRC15), NFκB2 emerges as a key link between myeloid inflammation and fibrotic remodeling (Fig. 4M-N). Interestingly, CD1C^+^ cDC2 were significantly decreased in ALCA compared to HG (Fig. 4D-F, Supplementary Fig. 6B), indicating weaker antigen presentation and immune regulation^31,32^. This reduction may undermine immune surveillance and exacerbate the chronic inflammatory milieu characteristic of ALCA.

### Lymphocyte Cell Heterogeneity in ALCA

ALCA hearts exhibited a significant rise in lymphocyte infiltration, especially among CD8^+^ T cells (Fig. 5A). These lymphocytes displayed upregulated expression of genes associated with TNF signaling, oxidative stress, and ECM organization (Fig. 5B-C, Supplementary Fig. 7A). Seven distinct lymphocyte subpopulations were identified, including CD8^+^ effector memory T cells (CD8^+^ Tem), CD69^+^ CD8^+^ T cells, MKI67^+^ CD8^+^ T cells, CD45^+^ fibroblast-like cells (FB-CD45), CD45^+^ endothelial-like cells (Endo-CD45), NK cells, and B cells (Fig. 5D-F, Supplementary Fig. 7B-C). Trajectory analysis suggested that CD8^+^ Tem could differentiate into other T cell subtypes, highlighting a dynamic response to ALCA pathology (Fig. 5G, Supplementary Fig. 7D). CD69^+^ CD8^+^ T cells were notably expanded in ALCA and highly expressed FOSB, CITED2, and TNF (Fig. 5H). Functional pathway analyses indicated involvement in ERK signaling, IL-17 signaling, and apoptosis-related genes (Fig. 5I–J), suggesting enhanced immune surveillance and cytotoxic capacity. TF analysis revealed elevated KLF6 activity in T cell subsets, modulating IRF2BP2 and RBPJ, underlining the pivotal role of KLF6 in T cell function in ALCA (Fig. 5K-M).

**Fig. 5.**
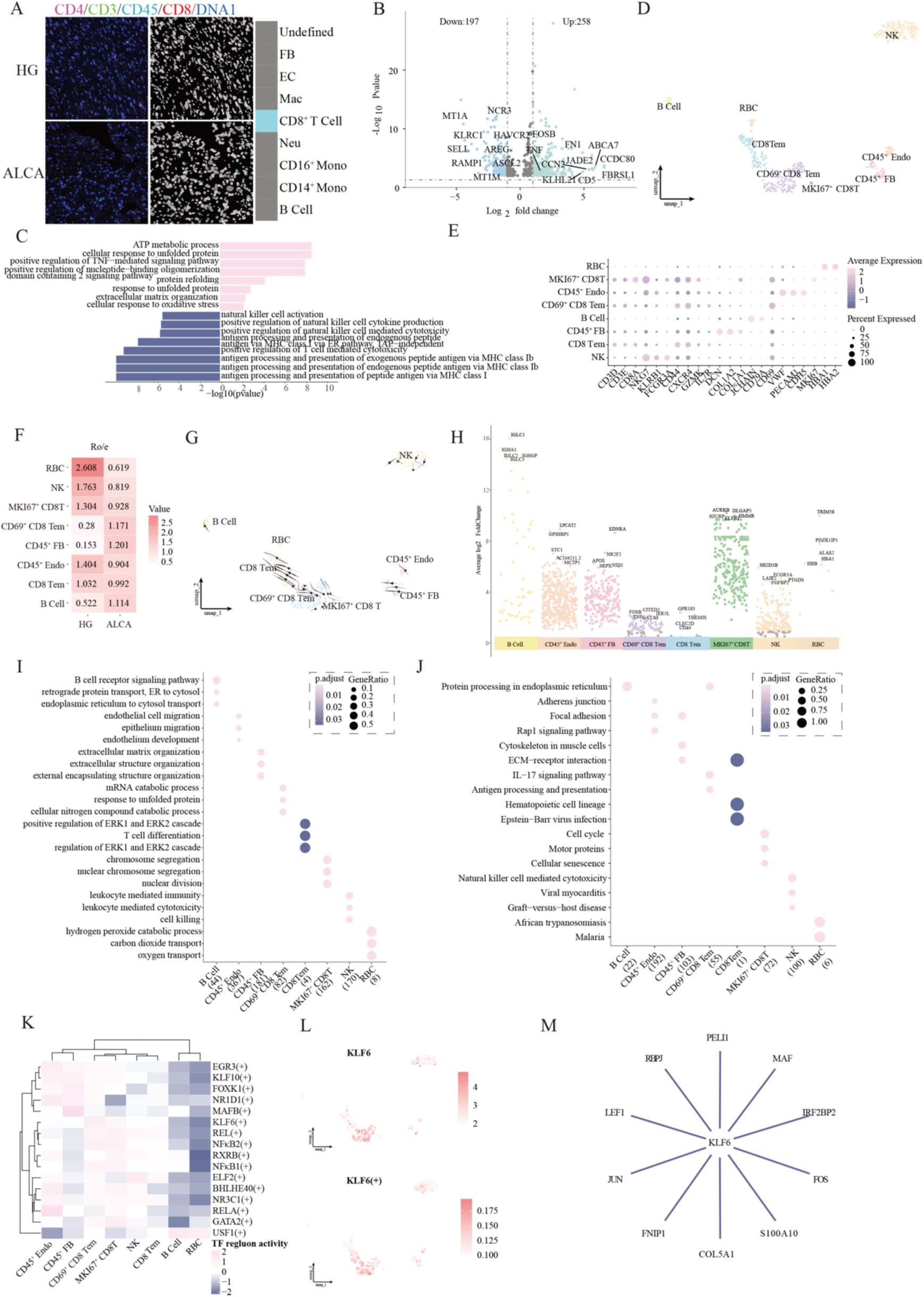
Characterization of lymphocyte cell in ALCA. **(A)** IMC images (left panel) and cell-type masks (right panel) with CD8^+^ T cells makers, including CD4 (magenta), CD3 (lime), CD45 (cyan), CD8 (red) and DNA1 (blue) in HG and ALCA hearts. **(B)** Volcano plot of DEGs in lymphocyte clusters. Blue dots represent significantly downregulated genes, while green dots indicate significantly upregulated genes. **(C)** Bar plot of GO enrichment for DEGs. **(D)** UMAP with eight lymphocyte subsets from scRNA-seq. **(E)** Dot plot of canonical markers in each lymphocyte cluster. **(F)** Ro/e scores summarizing cluster prevalence. **(G)** Velocity streamlines showing lymphocyte differentiation paths. **(H)** Top marker genes for various lymphocyte subtypes. **(I)** GO dot plots for lymphocyte subclusters. **(J)** KEGG dot plots for lymphocyte subclusters. **(K)** Heatmap of TF motif activity for the top five motifs in each lymphocyte subset. **(L)** UMAP shows expression (up) and activity (bottom) of KLF6. **(M)** Circos diagram linking lymphocyte signature genes (outer circle) to TF regulons (inner circle).

Conversely, NK cells were markedly reduced in ALCA tissues, typically expressing SH2D1B, FCGR3A, and PTGDs (Fig. 5H), which mediate leukocyte-driven immunity and NK cell–mediated cytotoxicity (Fig. 5I-J). This reduction may weaken innate immune responses, potentially exacerbating cardiac tissue injury^33^.

### Cellular Interactions and Intercellular Signaling in ALCA

Intercellular communication in ALCA markedly differs from HG, particularly in regions enriched for macrophages and activated fibroblasts (Fig. 6A). The interaction scores between these cell types were notably increased in ALCA, indicating a feedback loop that contributes to fibrotic remodeling (Fig. 6B).

**Fig. 6.**
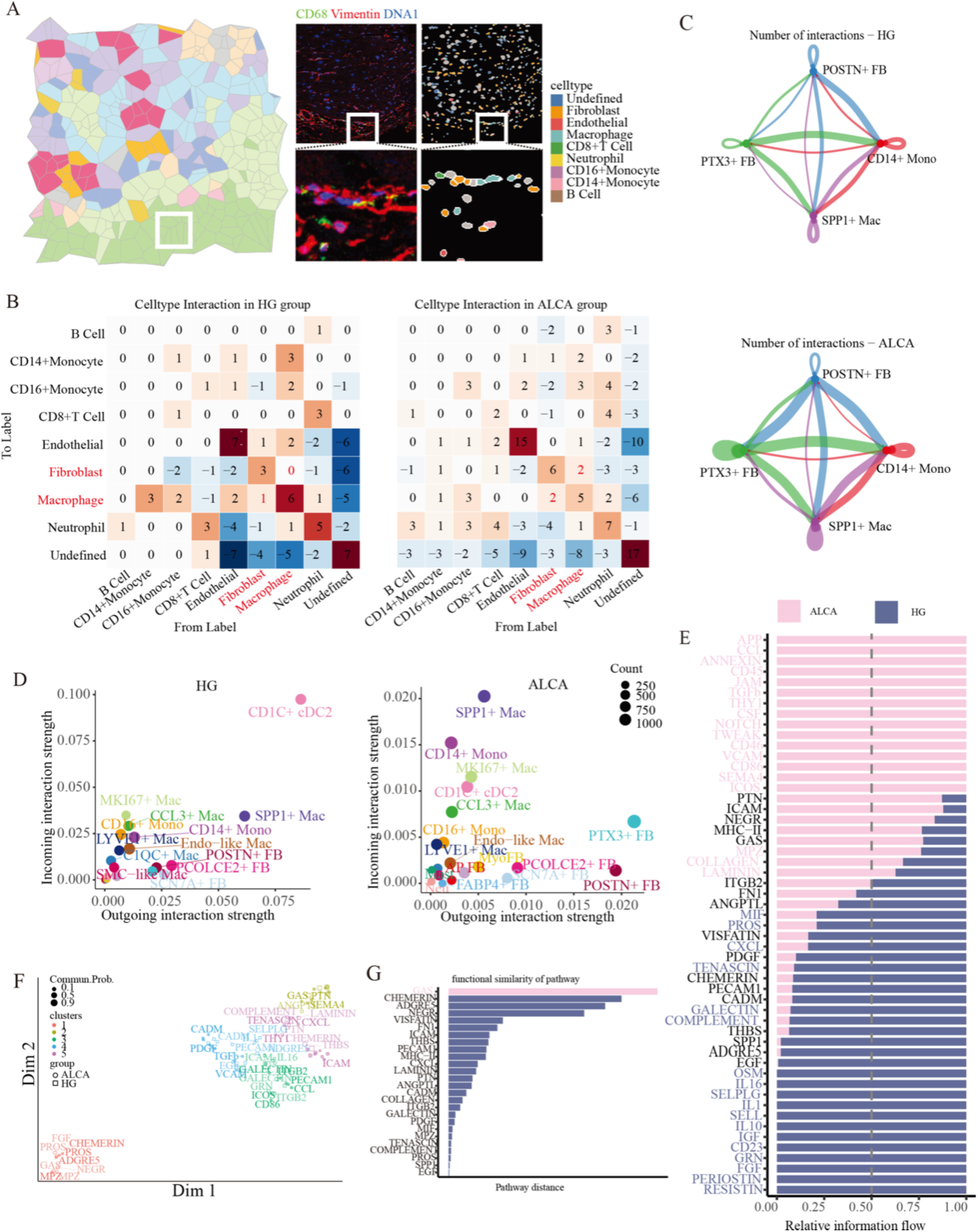
Fibroblast–Macrophage Interactions. **(A)** Voronoi diagrams (left panels), IMC images (middle panel), and cell-type masks (right panel) showing spatial distribution of macrophages (Mac) and fibroblasts (FB) in HG and ALCA. CD68 (lime), vimentin (red) and DNA1 (blue) were used to depict the structure of HG and ALCA heart. **(B)** Neighborhood analysis illustrating spatial interactions derived from IMC. Rows represent a centered cell type; columns are neighboring cell types. **(C)** Bar plot of major cell-type communications (POSTN^+^ FB, PTX3^+^ FB, CD14^+^ Mono, SPP1^+^ Mac) identified by CellChat. **(D)** Scatterplot of dominant senders and receivers, with dot size reflecting the number of predicted signaling links. **(E)** Ranked signaling pathways based on overall information flow differences between HG and ALCA. The top signaling pathways colored by pink are more enriched in ALCA, the bottom ones colored by blue were more enriched in the HG and the middle ones colored by black were non-significantly between the two groups. **(F)** Shared two-dimensional manifold clustering of signaling pathways by functional similarity. Circle and square symbols represent the signaling networks from ALCA and HG respectively. Each dot or square represents the communication network of one signaling pathway. Dot or square size is proportional to the total communication probability. Different colors represent different groups of signaling pathways. **(G)** Functional divergence of signaling pathways between HG and ALCA, measured by Euclidean distance. Larger distances indicate greater functional divergence between the communication networks of the two datasets.

A cell interaction matrix encompassing CD14^+^ Mono, SPP1^+^ Mac, PTX3^+^ FB, and POSTN^+^ FB highlighted a significantly increase in crosstalk within ALCA (Fig. 6C, Supplementary Fig. 8A). SPP1^+^ Mac, known for their pro-angiogenic and pro-fibrotic roles^34,35^, may directly interact with PTX3^+^ FB to drive cardiac fibrosis. Meanwhile, CD1C^+^ cDC2 were decreased, shifting the overall signaling balance toward macrophage–fibroblast pathways in ALCA (Fig. 6D).

Comparative analysis of ALCA and HG showed a significant elevation of multiple signaling pathways (PTN, ICAM, ITGB2, NEGR, FN1), as well as ALCA-specific networks (APP, CCL, TGF-β, CSF) (Fig. 6E). Joint manifold learning clustered these pathways, showing substantial functional shifts (Fig. 6F). Notably, GAS6 exhibited the greatest divergence between ALCA and HG (Fig. 6G).

CellChat analysis underscored pivotal crosstalk between macrophages and fibroblasts, particularly along the SPP1^+^ Mac–PTX3^+^ FB axis. Specifically, SPP1^+^ Mac–PTX3^+^ FB interactions (TGF-β1–TGF-βR1/2, NAMPT–INSR, LGALS9–CD44) were linked to enhanced myofibroblast differentiation and ECM deposition (Fig. 7A, Supplementary Fig. 8B). In turn, PTX3^+^ FB supported SPP1^+^ Mac polarization via GAS6–MERTK, CSF1–CSF1R, and APP–CD74 (Fig. 7 B-C, Supplementary Fig. 8C and F). This reciprocal signaling loop may amplify immune-driven fibrosis and drive disease progression. PTX3^+^ FB and SPP1^+^ Mac also facilitated CD14^+^ Mono adhesion and differentiation via pathways like CSF1–CSF1R (Supplementary Fig. 8D-E). Although POSTN^+^ FB and CD14^+^ Mono also contributed to the ALCA microenvironment, the SPP1^+^ Mac–PTX3^+^ FB axis emerged as a primary driver of chronic inflammation and fibrotic remodeling in ALCA.

**Fig. 7.**
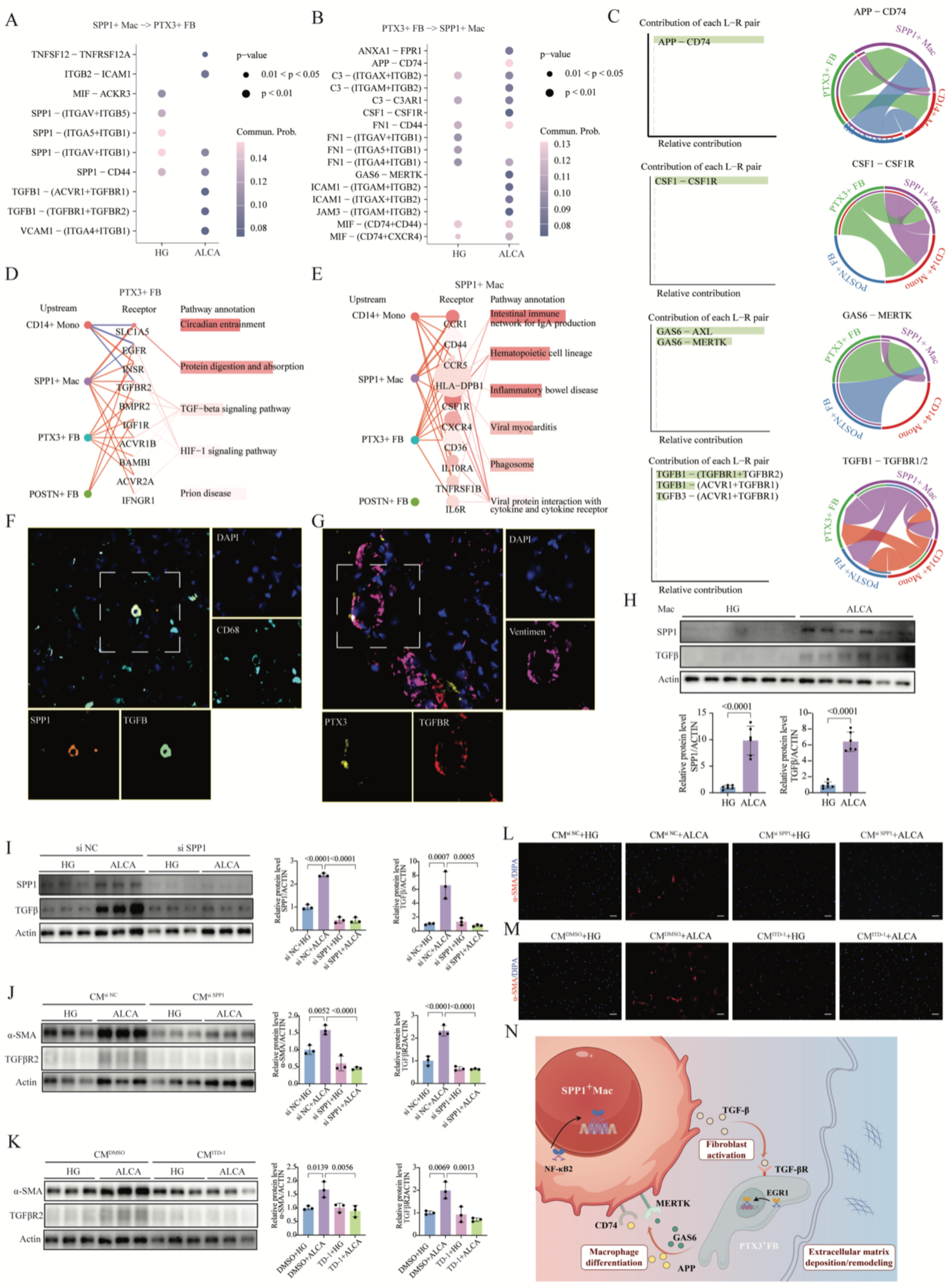
Key Ligand–Receptor Interactions Between Fibroblasts and Macrophages. (A-B) Representative ligand-receptor pairs from SPP1^+^ Mac to PTX3^+^ FB, and PTX3^+^ FB to SPP1^+^ Mac in HG and ALCA. Circle size indicates significance, color indicates interaction strength. **(C)** Bar graph (left) of ligand–receptor contributions in PTX3^+^ FB and SPP1^+^ Mac, circle plot (right) of top ligand–receptor networks. Arrow root indicates the ligand-expressing cell type, arrow tip is the receiving cell. **(D-E)** Pathway-mediated communication chains for PTX3^+^ FB and SPP1^+^ Mac, highlighting significantly upregulated pathways compared to other cardiac cells. Line widths indicate overall interaction intensity (Upstream → Receptor), while dot size and color in the receptor column reflect log₂ fold change and −log₁₀(p-value). **(F-G)** Representative mIHC images of fFFPE heart tissue from HG and ALCA, stained for CD68 (azure), TGFβ (green), SPP1 (orange), PTX3 (yellow), TGFβR (red), Vimentin (purple), and DAPI (blue). **(H)** Immunoblotting analysis for SPP1 and TGFβ1 protein expression in macrophages under stimulation with ALCA patient serum. **(I)** Immunoblotting analysis for SPP1 and TGFβ1 protein expression in macrophages with si SPP1 under stimulation with ALCA patient serum. **(J)** Immunoblotting analysis for TGFβR and α-SMA protein expression in fibroblasts co-culture with conditioned medium from macrophages. **(K)** Immunoblotting analysis for TGF-βR and α-SMA protein expression in fibroblasts co-culture with conditioned medium from macrophages following TGF-βR2 inhibition (ITD-1). **(L-M)** Representative immunofluorescence images of fibroblasts co-culture with conditioned medium from macrophages, stained for α-SMA (red), and DAPI (blue). **(N)** Schematic representation of the proposed ALCA mechanism.

Network centrality and CommPath analyses spotlighted PTX3^+^ FB and SPP1^+^ Mac as primary signaling hubs in ALCA. PTX3^+^ FB secretes CSF1 and GAS6, which engage CSF1R and MERTK on SPP1^+^ Mac, promoting macrophage recruitment and polarization. Conversely, PTX3^+^ FB responds to TGF-β1 from SPP1^+^ Mac via TGF-βR1/2, driving myofibroblast transformation (Fig. 7D-E, Supplementary Fig. 8G-H). This bidirectional loop not only propels fibrotic remodeling but also orchestrates a pro-inflammatory microenvironment (Fig. 7D-E). Although POSTN^+^ FB and CD14^+^ Mono both support the overall fibrotic milieu, SPP1^+^ Mac–PTX3^+^ FB interactions dominate. In situ multiplex immunohistochemistry (mIHC) verified these computational findings by revealing higher TGF-β1 in SPP1^+^ Mac and enhanced TGF-βR2 in PTX3^+^ FB (Fig. 7F-G).

Furthermore, an in vitro co-culture model using primary macrophage and fibroblast demonstrated that macrophages stimulated by ALCA patient serum displayed elevated SPP1 and TGF-β1 expression (Fig. 7H). Deletion of SPP1 significantly reduce TGF-β1 expression (Fig. 7I). Conditioned medium (CM) derived from macrophage induced notable upregulation of TGF-βR2 and α-SMA in fibroblasts, indicative of myofibroblast transformation (Fig. 7J). Inhibition of either SPP1 or TGF-βR2 partially reversed these fibrotic markers (Fig. 7K). Immunofluorescence confirmed increased α-SMA expression in fibroblasts exposed to ALCA macrophage-CM, which decreased following inhibition of SPP1 and TGF-βR2 (Fig. 7L-M). Collectively, these data underscore the central role of SPP1^+^ Mac–PTX3^+^ FB crosstalk in orchestrating pathogenic inflammation and fibrosis in ALCA (Fig. 7N).

## Discussion

This study presents the first comprehensive single-cell and spatial landscape of ALCA, illustrating how diverse cell populations drive fibrosis, immune dysregulation, and endothelial dysfunction. By mapping intercellular interactions at high resolution, we have pinpointed molecular targets and pathways that may mitigate the deleterious consequences of amyloid deposition.

Our study sheds light on the significant fibrotic remodeling in ALCA, aligning with prior reports indicating myocardial fibrosis in ALCA patients, which did not spatially overlap with amyloid and contributed to elevated extracellular volume^36^. Importantly, our findings suggest that fibrosis in ALCA is an active, dynamic process mediated by TGF-β-driven myofibroblast activation rather than a passive accumulation of static scar tissue. A prominent discovery is the identification of distinct fibroblast subtypes critical for fibrosis. Early-stage PTX3^+^ FB transition into pro-fibrotic POSTN^+^ FB through transcriptional programs regulated by EGR1, KLFs, and NFκB1/2. Notably, PTX3^+^ FBs exhibit high expression of genes involved in immune responses, ECM remodeling, and misfolded protein processing. Therapeutic strategies targeting PTX3^+^ FB may curb ECM deposition and delay the transition to a fully fibrotic state, consistent with the established roles of PTX3 in tissue repair and immune regulation^37–39^.

Endothelial dysfunction emerged as another crucial facet of ALCA pathology^40,41^. We identified distinct endothelial subtypes, notably inflammatory venous EC and capillary-like immune EC, which produce chemokines (CXCL1, CXCL3, CCL2) that likely sustain chronic immune infiltration^42^. Concurrent depletion of functional capillary EC and smooth muscle-like EC populations further exacerbates vascular dysfunction, suggesting compromised myocardial perfusion^43^. Therapeutically targeting endothelial dysfunction and its associated inflammatory chemokines may offer strategies to preserve cardiac function in ALCA patients.

Our study also provides compelling evidence for macrophages as primary drivers of fibrosis and chronic inflammation. SPP1^+^ Mac plays a particularly pivotal role, promoting fibroblast differentiation and ECM deposition through direct interactions involving TGF-Β1–TGF-ΒR1/2 and GAS6–MERTK pathways. CCL3^+^ Mac further perpetuates inflammation, reinforcing the pathogenic milieu. The experimental validation of SPP1^+^ Mac–PTX3^+^ FB signaling axis via co-culture experiments further supports their central role. Moreover, we highlight NFκB2 signaling as a critical regulator integrating immune responses and fibrosis, emphasizing its potential as a therapeutic target to rebalance inflammatory homeostasis^44^.

Additionally, we highlight a crucial reciprocal signaling loop between PTX3^+^ fibroblasts and SPP1^+^ macrophages. PTX3^+^ FB not only facilitate differentiation of monocytes to pro-fibrotic macrophages through CSF1–CSF1R and APP–CD74 pathways but also respond to macrophage-derived TGF-β, driving their myofibroblast transformation. Intervening in this feedback loop could effectively reduce fibrosis and chronic inflammation, potentially halting ALCA progression. The expansion of CD69^+^ CD8^+^ T cells indicate enhanced cytotoxic surveillance^45^, whereas the marked reduction of NK cells likely impairs the capacity to eliminate damaged cells^33^. Restoring NK cell function or rebalancing cytotoxic T-cell responses may help alleviate chronic inflammation and limit myocardial injury.

## Conclusion

This study provides a detailed atlas of the cellular landscape and intercellular signaling in ALCA, revealing how distinct cell populations, particularly PTX3^+^ FB and SPP1^+^ Mac, orchestrate fibrosis, immune dysregulation, and endothelial dysfunction. we propose novel therapeutic targets that could significantly enhance the clinical management of cardiac amyloidosis and offer new directions for translational research.

## Limitations of the study

Despite the scope of this investigation, certain constraints remain. First, the relatively small sample size may restrict the broader applicability of our observations. Second, the absence of robust ALCA animal models limited our ability to extensively validate the proposed mechanisms in vivo. Consequently, future studies leveraging larger patient cohorts, well-designed clinical trials, and improved experimental models will be crucial for validating and extending these findings. Nonetheless, our single-cell atlas provides a foundation for novel, cell-targeted therapeutic strategies in ALCA.

## Methods

All experiments were performed according to standard protocols. Full methodological details are provided in the Supplementary Materials

## Data availability

The data and code that support the findings of this study are available from the corresponding author upon request.

## Acknowledgements

We extend our sincere gratitude to the donors and their families for providing the tissue samples that made this research possible. We also thank the Cardiovascular Biobank and Resource Center at the Second Affiliated Hospital, Zhejiang University School of Medicine for their invaluable support. In addition, we acknowledge Hangzhou Kaitai Biotechnology Co., Ltd. (Hangzhou, China) and Infinity Scope Spatial Multi-Omics Biotechnology Co., Ltd. for their contributions. Figure 7N was created using Figdraw (www.figdraw.com).

## Author contributions

Conceptualization: Q.M, Z.W, N.Z, and X.P. Methodology: Q.M, Z.W, and J.Z. Writing—original draft: Q.M and Z.W. Writing—review and editing: Q.M, Z., J.Z, X.L, J.X, X.P and N.Z. Funding acquisition: N.Z. Resources: X.P and Q.M. Supervision: X.P and N.Z. Data curation: J.Z, X.L.

## Reference

1. Kittleson MM, Maurer MS, Ambardekar AV, Bullock-Palmer RP, Chang PP, Eisen HJ, Nair AP, Nativi-Nicolau J, Ruberg FL, American Heart Association Heart F, et al. Cardiac Amyloidosis: Evolving Diagnosis and Management: A Scientific Statement From the American Heart Association. Circulation. 2020;142:e7–e22. doi: 10.1161/CIR.0000000000000792

2. Sanchorawala V. Systemic Light Chain Amyloidosis. The New England journal of medicine. 2024;390:2295–2307. doi: 10.1056/NEJMra2304088

3. Smiley DA, Einstein AJ. From the Bone Marrow to the Heart: Cardiac Recovery in AL Amyloidosis. JACC Asia. 2025;5:85–87. doi: 10.1016/j.jacasi.2024.12.001

4. Ravichandran S, Lachmann HJ, Wechalekar AD. Epidemiologic and Survival Trends in Amyloidosis, 1987-2019. The New England journal of medicine. 2020;382:1567–1568. doi: 10.1056/NEJMc1917321

5. Alexander KM, Orav J, Singh A, Jacob SA, Menon A, Padera RF, Kijewski MF, Liao R, Di Carli MF, Laubach JP, et al. Geographic Disparities in Reported US Amyloidosis Mortality From 1979 to 2015: Potential Underdetection of Cardiac Amyloidosis. JAMA Cardiol. 2018;3:865–870. doi: 10.1001/jamacardio.2018.2093

6. Griffin JM, Rosenblum H, Maurer MS. Pathophysiology and Therapeutic Approaches to Cardiac Amyloidosis. Circulation research. 2021;128:1554–1575. doi: 10.1161/CIRCRESAHA.121.318187

7. Zhang D, Wen Q, Zhang R, Kou K, Lin M, Zhang S, Yang J, Shi H, Yang Y, Tan X, et al. From Cell to Gene: Deciphering the Mechanism of Heart Failure With Single-Cell Sequencing. Adv Sci (Weinh). 2024;11:e2308900. doi: 10.1002/advs.202308900

8. Litvinukova M, Talavera-Lopez C, Maatz H, Reichart D, Worth CL, Lindberg EL, Kanda M, Polanski K, Heinig M, Lee M, et al. Cells of the adult human heart. Nature. 2020;588:466–472. doi: 10.1038/s41586-020-2797-4

9. Jiang Z, Kang Q, Qian H, Xu Z, Tong H, Yang J, Li L, Li R, Li G, Chen F, et al. Revealing the crucial roles of suppressive immune microenvironment in cardiac myxoma progression. Signal Transduct Target Ther. 2024;9:193. doi: 10.1038/s41392-024-01912-2

10. Farah EN, Hu RK, Kern C, Zhang Q, Lu TY, Ma Q, Tran S, Zhang B, Carlin D, Monell A, et al. Spatially organized cellular communities form the developing human heart. Nature. 2024;627:854–864. doi: 10.1038/s41586-024-07171-z

11. Hickey JW, Neumann EK, Radtke AJ, Camarillo JM, Beuschel RT, Albanese A, McDonough E, Hatler J, Wiblin AE, Fisher J, et al. Spatial mapping of protein composition and tissue organization: a primer for multiplexed antibody-based imaging. Nat Methods. 2022;19:284–295. doi: 10.1038/s41592-021-01316-y

12. Moffitt JR, Lundberg E, Heyn H. The emerging landscape of spatial profiling technologies. Nat Rev Genet. 2022;23:741–759. doi: 10.1038/s41576-022-00515-3

13. Zhu H, Galdos FX, Lee D, Waliany S, Huang YV, Ryan J, Dang K, Neal JW, Wakelee HA, Reddy SA, et al. Identification of Pathogenic Immune Cell Subsets Associated With Checkpoint Inhibitor-Induced Myocarditis. Circulation. 2022;146:316–335. doi: 10.1161/CIRCULATIONAHA.121.056730

14. Zhang K, Wang Y, Chen S, Mao J, Jin Y, Ye H, Zhang Y, Liu X, Gong C, Cheng X, et al. TREM2(hi) resident macrophages protect the septic heart by maintaining cardiomyocyte homeostasis. Nat Metab. 2023;5:129–146. doi: 10.1038/s42255-022-00715-5

15. Servaas NH, Hiddingh S, Chouri E, Wichers CGK, Affandi AJ, Ottria A, Bekker CPJ, Cossu M, Silva-Cardoso SC, van der Kroef M, et al. Nuclear Receptor Subfamily 4A Signaling as a Key Disease Pathway of CD1c+ Dendritic Cell Dysregulation in Systemic Sclerosis. Arthritis Rheumatol. 2023;75:279–292. doi: 10.1002/art.42319

16. Liu RJ, Niu XL, Yuan JP, Chen HD, Gao XH, Qi RQ. DnaJA4 is involved in responses to hyperthermia by regulating the expression of F-actin in HaCaT cells. Chin Med J (Engl). 2020;134:456–462. doi: 10.1097/CM9.0000000000001064

17. von Scheidt M, Zhao Y, de Aguiar Vallim TQ, Che N, Wierer M, Seldin MM, Franzen O, Kurt Z, Pang S, Bongiovanni D, et al. Transcription Factor MAFF (MAF Basic Leucine Zipper Transcription Factor F) Regulates an Atherosclerosis Relevant Network Connecting Inflammation and Cholesterol Metabolism. Circulation. 2021;143:1809–1823. doi: 10.1161/CIRCULATIONAHA.120.050186

18. Meng Q, Yang B, Qiao Y, Wu Y, Chen J, Lin X, Molkentin JD. Genetic and Pharmacologic Inhibition of JAK1/2 Antagonizes Cardiac Fibrosis. Circulation. 2024;150:899–901. doi: 10.1161/CIRCULATIONAHA.124.070340

19. Alexanian M, Przytycki PF, Micheletti R, Padmanabhan A, Ye L, Travers JG, Gonzalez-Teran B, Silva AC, Duan Q, Ranade SS, et al. A transcriptional switch governs fibroblast activation in heart disease. Nature. 2021;595:438–443. doi: 10.1038/s41586-021-03674-1

20. Poulsen PC, Schrolkamp M, Bagwan N, Leurs U, Humphries ESA, Bomholzt SH, Nielsen MS, Bentzen BH, Olsen JV, Lundby A. Quantitative proteomics characterization of acutely isolated primary adult rat cardiomyocytes and fibroblasts. J Mol Cell Cardiol. 2020;143:63–70. doi: 10.1016/j.yjmcc.2020.04.021

21. Gautheron J, Elsayed S, Pistorio V, Lockhart S, Zammouri J, Auclair M, Koulman A, Meadows SR, Lhomme M, Ponnaiah M, et al. ADH1B, the adipocyte-enriched alcohol dehydrogenase, plays an essential, cell-autonomous role in human adipogenesis. Proceedings of the National Academy of Sciences of the United States of America. 2024;121:e2319301121. doi: 10.1073/pnas.2319301121

22. Wang X, Guo Y, Gao Y, Ren C, Huang Z, Liu B, Li X, Chang L, Shen K, Ding H, et al. Feasibility of (68)Ga-Labeled Fibroblast Activation Protein Inhibitor PET/CT in Light-Chain Cardiac Amyloidosis. JACC Cardiovascular imaging. 2022;15:1960–1970. doi: 10.1016/j.jcmg.2022.06.004

23. Hahn VS, Yanek LR, Vaishnav J, Ying W, Vaidya D, Lee YZJ, Riley SJ, Subramanya V, Brown EE, Hopkins CD, et al. Endomyocardial Biopsy Characterization of Heart Failure With Preserved Ejection Fraction and Prevalence of Cardiac Amyloidosis. JACC Heart Fail. 2020;8:712–724. doi: 10.1016/j.jchf.2020.04.007

24. Gama F, Rosmini S, Bandula S, Patel KP, Massa P, Tobon-Gomez C, Ecke K, Stroud T, Condron M, Thornton GD, et al. Extracellular Volume Fraction by Computed Tomography Predicts Long-Term Prognosis Among Patients With Cardiac Amyloidosis. JACC Cardiovascular imaging. 2022;15:2082–2094. doi: 10.1016/j.jcmg.2022.08.006

25. Cook ME, Jarjour NN, Lin CC, Edelson BT. Transcription Factor Bhlhe40 in Immunity and Autoimmunity. Trends Immunol. 2020;41:1023–1036. doi: 10.1016/j.it.2020.09.002

26. Kalucka J, de Rooij L, Goveia J, Rohlenova K, Dumas SJ, Meta E, Conchinha NV, Taverna F, Teuwen LA, Veys K, et al. Single-Cell Transcriptome Atlas of Murine Endothelial Cells. Cell. 2020;180:764–779 e720. doi: 10.1016/j.cell.2020.01.015

27. Qi J, Sun H, Zhang Y, Wang Z, Xun Z, Li Z, Ding X, Bao R, Hong L, Jia W, et al. Single-cell and spatial analysis reveal interaction of FAP(+) fibroblasts and SPP1(+) macrophages in colorectal cancer. Nature communications. 2022;13:1742. doi: 10.1038/s41467-022-29366-6

28. Hulsmans M, Schloss MJ, Lee IH, Bapat A, Iwamoto Y, Vinegoni C, Paccalet A, Yamazoe M, Grune J, Pabel S, et al. Recruited macrophages elicit atrial fibrillation. Science. 2023;381:231–239. doi: 10.1126/science.abq3061

29. Spadaro O, Youm Y, Shchukina I, Ryu S, Sidorov S, Ravussin A, Nguyen K, Aladyeva E, Predeus AN, Smith SR, et al. Caloric restriction in humans reveals immunometabolic regulators of health span. Science. 2022;375:671–677. doi: 10.1126/science.abg7292

30. Caronni N, La Terza F, Vittoria FM, Barbiera G, Mezzanzanica L, Cuzzola V, Barresi S, Pellegatta M, Canevazzi P, Dunsmore G, et al. IL-1beta(+) macrophages fuel pathogenic inflammation in pancreatic cancer. Nature. 2023;623:415–422. doi: 10.1038/s41586-023-06685-2

31. Tiberio L, Del Prete A, Schioppa T, Sozio F, Bosisio D, Sozzani S. Chemokine and chemotactic signals in dendritic cell migration. Cell Mol Immunol. 2018;15:346–352. doi: 10.1038/s41423-018-0005-3

32. Del Prete A, Salvi V, Soriani A, Laffranchi M, Sozio F, Bosisio D, Sozzani S. Dendritic cell subsets in cancer immunity and tumor antigen sensing. Cell Mol Immunol. 2023;20:432–447. doi: 10.1038/s41423-023-00990-6

33. Rodriguez-Iturbe B, Pons H, Johnson RJ. Role of the Immune System in Hypertension. Physiol Rev. 2017;97:1127–1164. doi: 10.1152/physrev.00031.2016

34. Dahlgren MW, Molofsky AB. Adventitial Cuffs: Regional Hubs for Tissue Immunity. Trends Immunol. 2019;40:877–887. doi: 10.1016/j.it.2019.08.002

35. Gao Y, Li J, Cheng W, Diao T, Liu H, Bo Y, Liu C, Zhou W, Chen M, Zhang Y, et al. Cross-tissue human fibroblast atlas reveals myofibroblast subtypes with distinct roles in immune modulation. Cancer Cell. 2024;42:1764–1783 e1710. doi: 10.1016/j.ccell.2024.08.020

36. Pucci A, Aimo A, Musetti V, Barison A, Vergaro G, Genovesi D, Giorgetti A, Masotti S, Arzilli C, Prontera C, et al. Amyloid Deposits and Fibrosis on Left Ventricular Endomyocardial Biopsy Correlate With Extracellular Volume in Cardiac Amyloidosis. J Am Heart Assoc. 2021;10:e020358. doi: 10.1161/JAHA.120.020358

37. Bonavita E, Gentile S, Rubino M, Maina V, Papait R, Kunderfranco P, Greco C, Feruglio F, Molgora M, Laface I, et al. PTX3 is an extrinsic oncosuppressor regulating complement-dependent inflammation in cancer. Cell. 2015;160:700–714. doi: 10.1016/j.cell.2015.01.004

38. Garlanda C, Bottazzi B, Magrini E, Inforzato A, Mantovani A. PTX3, a Humoral Pattern Recognition Molecule, in Innate Immunity, Tissue Repair, and Cancer. Physiol Rev. 2018;98:623–639. doi: 10.1152/physrev.00016.2017

39. Salio M, Chimenti S, De Angelis N, Molla F, Maina V, Nebuloni M, Pasqualini F, Latini R, Garlanda C, Mantovani A. Cardioprotective function of the long pentraxin PTX3 in acute myocardial infarction. Circulation. 2008;117:1055–1064. doi: 10.1161/CIRCULATIONAHA.107.749234

40. Chacko L, Kotecha T, Ioannou A, Patel N, Martinez-Naharro A, Razvi Y, Patel R, Massa P, Venneri L, Brown J, et al. Myocardial perfusion in cardiac amyloidosis. Eur J Heart Fail. 2024;26:598–609. doi: 10.1002/ejhf.3137

41. Siddiqi OK, Ruberg FL. Cardiac amyloidosis: An update on pathophysiology, diagnosis, and treatment. Trends Cardiovasc Med. 2018;28:10–21. doi: 10.1016/j.tcm.2017.07.004

42. Drummer Ct, Saaoud F, Shao Y, Sun Y, Xu K, Lu Y, Ni D, Atar D, Jiang X, Wang H, et al. Trained Immunity and Reactivity of Macrophages and Endothelial Cells. Arterioscler Thromb Vasc Biol. 2021;41:1032–1046. doi: 10.1161/ATVBAHA.120.315452

43. Joo HJ, Choi DK, Lim JS, Park JS, Lee SH, Song S, Shin JH, Lim DS, Kim I, Hwang KC, et al. ROCK suppression promotes differentiation and expansion of endothelial cells from embryonic stem cell-derived Flk1(+) mesodermal precursor cells. Blood. 2012;120:2733–2744. doi: 10.1182/blood-2012-04-421610

44. Sun SC. The non-canonical NF-kappaB pathway in immunity and inflammation. Nat Rev Immunol. 2017;17:545–558. doi: 10.1038/nri.2017.52

45. Taylor RP. T cells reinforce NK cell-mediated ADCC. Blood. 2024;143:1786–1787. doi: 10.1182/blood.2024024444

